# Leaf traits drive differences in biomass partitioning among major plant functional types

**DOI:** 10.1101/025361

**Authors:** Remko A. Duursma, Daniel S. Falster

## Abstract

1. The partitioning of biomass into leaves and stems is one of the most uncertain and influential components of global vegetation models (GVMs). Although GVMs typically assume that the major woody plant functional types (PFTs) differ in biomass partitioning, empirical studies have not been able to justify these differences. Here we test for differences between PFTs in partitioning of biomass between leaves and stems.
2. We use the recently published Biomass And Allometry Database (BAAD), a large database including observations for individual plants. The database covers the global climate space, allowing us to test for direct climate effects in addition to PFT.
3. The leaf mass fraction (LMF, leaf / total aboveground biomass) varied strongly between PFTs (as defined by deciduous vs. evergreen and gymnosperm vs. angiosperm). We found that LMF, once corrected for plant height, was proportional to leaf mass per area across PFTs. As a result, the PFTs did not differ in the amount of leaf area supported per unit above ground biomass. We found only weak and inconsistent effects of climate on biomass partitioning.
4. Combined, these results uncover fundamental rules in how plants are constructed and allow for systematic benchmarking of biomass partitioning routines in GVMs.

## Introduction

The partitioning of forest biomass among leaves and stems strongly influences the productivity and carbon cycle of the world’s vegetation (Ise *et al.*, 2010; De Kauwe *et al.*, 2014; Friend *et al.*, 2014). Biomass stored in woody stems has a long residence time (Luyssaert *et al.*, 2008), whereas leaf biomass turns over quickly, entering the soil carbon cycle where the majority of carbon is released back to the atmosphere (Ryan & Law, 2005). Globally, forests store approximately 360Pg of carbon in living biomass (Pan *et al.*, 2011), equivalent to almost 40 years of current anthropogenic CO_2_ emissions (Friedlingstein *et al.*, 2014). Reducing uncertainties about biomass partitioning and carbon residence times in GVMs is therefore a priority for understanding the effects of climate and other environmental change on the global carbon cycle (Friend *et al.*, 2014; De Kauwe *et al.*, 2014; Negrón-Jurez *et al.*, 2015).

Perhaps the biggest challenge for GVMs is to capture the combined responses of the more than 250,000 plant species comprising the world’s vegetation. While most plants have the same basic resource requirements and physiological function, large differences exist among species in the amount of energy invested in different tissues (leaves, stems, roots). The approach taken by most GVMs for dealing with this functional diversity is to consider only a few archetypal plant functional types (PFTs) (Harrison *et al.*, 2010; Wullschleger *et al.*, 2014), assumed to differ in key physiological attributes. While GVMs assume or predict differences in biomass partitioning between PFTs (Notes S1), these differences are poorly constrained, due to limited available data. Moreover, there is little consensus on how biomass partitioning and allocation (see Methods for terminology) should be modelled in GVMs (Franklin *et al.*, 2012; De Kauwe *et al.*, 2014; Friend *et al.*, 2014). These shortcomings largely reflect the lack of suitable datasets of global scope with which models can be tested, constrained and compared (Wolf *et al.*, 2011).

In this paper we are primarily interested in the distribution of biomass (‘partitioning’) between leaves and woody stems, an important component of the residence time of carbon in ecosystems (Friend *et al.*, 2014). Previous work based on either whole stands (O’Neill & DeAngelis, 1981; Enquist & Niklas, 2002; Reich *et al.*, 2014) and or a mix on stand– and individual-based measurements (Poorter *et al.*, 2012, 2015), reported differences between angiosperms and gymnosperms in the amount of leaf biomass per unit above-ground biomass (the ‘leaf mass fraction’, LMF). Poorter *et al.* (2012) and Enquist & Niklas (2002) attributed higher LMF in gymnosperms to longer leaf lifespan (LL) compared to typical angiosperms, but this begs the question whether differences in LMF are also apparent between deciduous and evergreen functional types within angiosperms. It is also unknown whether PFTs with higher leaf biomass also have higher total leaf area, which is relevant because leaf area drives total light interception and thus productivity. Some oft-cited studies have also assumed that gymnosperms carry more leaf area than angiosperms (Chabot & Hicks, 1982; Bond, 1989).

Little is known about global-scale patterns in LMF and LAR in relation to climate. It can be expected that biomass partitioning is correlated with precipitation or mean annual temperature because smaller-scale comparisons have shown responsiveness of biomass partitioning to environmental drivers (Berninger & Nikinmaa, 1994; Callaway *et al.*, 1994; Delucia *et al.*, 2000; Poyatos *et al.*, 2007). Moreover, a recent study by Reich *et al.* (2014) demonstrated that stand-scale biomass partitioning was related to mean annual temperature across diverse forest stands. Indeed, many GVMs assume that climate affects biomass partitioning within a given PFT, often through soil water stress or other abiotic stress factors (see Notes S1). Again, however, these models do not agree on the degree of plasticity in biomass partitioning, or which climate variables it should respond to.

Despite considerable advances in theory underlying allometric scaling in plants (Enquist, 2003; West *et al.*, 1999; Savage *et al.*, 2010) we do not yet have a clear understanding of potential differences between PFTs in terms of biomass partitioning. Instead, most previous work has focussed on understanding size-related shifts in biomass partitioning, as governed by constraints including hydraulic supply and mechanical stability (Savage *et al.*, 2010). Rather than to advance a specific model of biomass partitioning or allocation, our view is that a broader evidence base is needed first to elucidate patterns between PFTs. This should enable those building GVMs to refine algorithms and parameter values to more closely match the observations. Here, we use a unique, new database (Falster *et al.*, 2015) (Fig. 1) to establish general rules on how biomass partitioning differs among three dominant woody PFTs across the globe: evergreen gymnosperms, evergreen angiosperms, and deciduous angiosperms. A recent compilation of plant biomass data (Poorter *et al.*, 2015) speculated that differences between gymnosperms and angiosperms in distribution of biomass between leaves and stems is related to differences in leaf mass per area. Here we can directly test this hypothesis as our database, unlike those of Enquist & Niklas (2002), Reich *et al.* (2014), and Poorter *et al.* (2015), includes many observations of leaf area as well as leaf mass measured on the same plants.

**Fig. 1.**
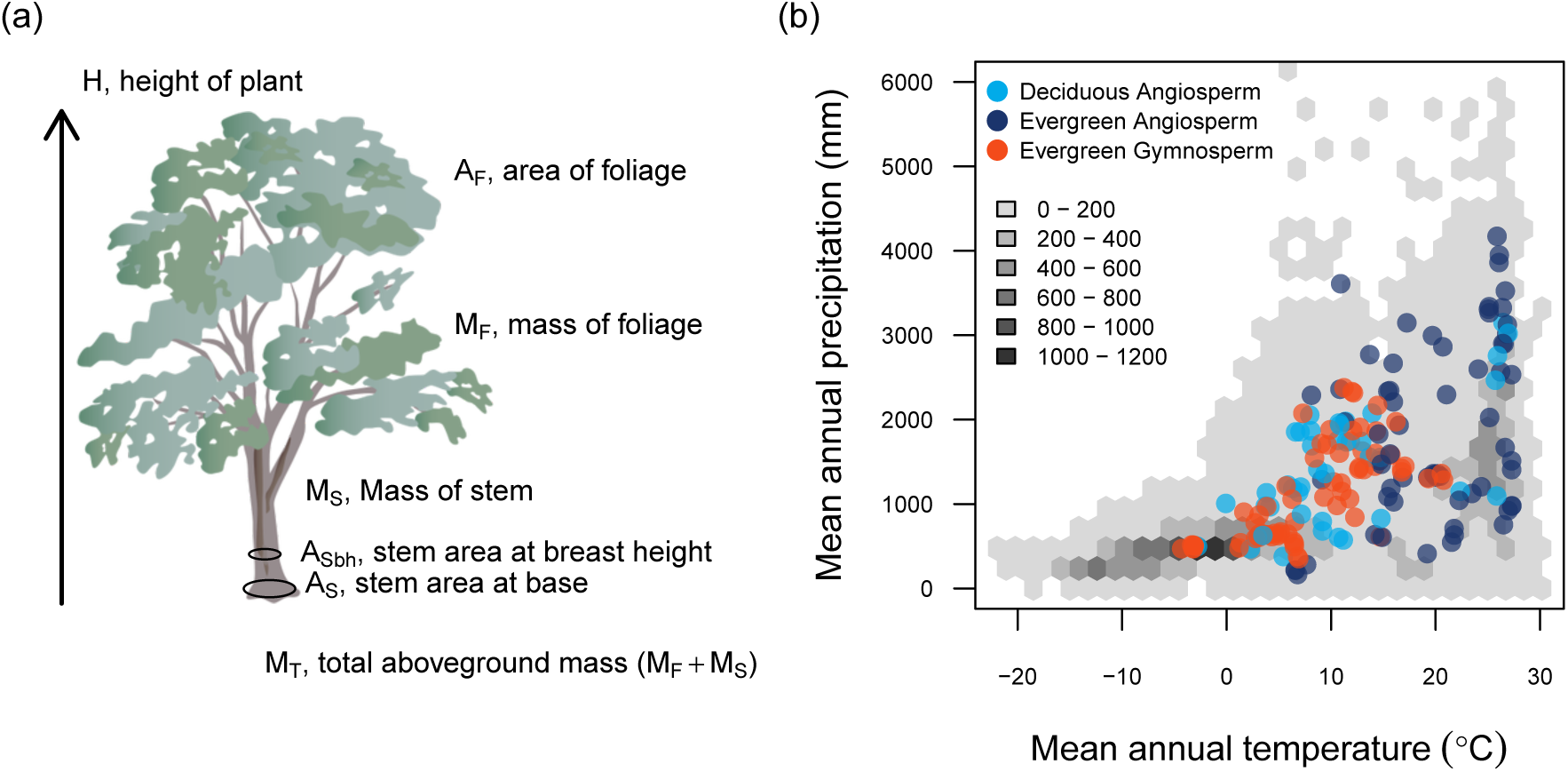
Overview of the data. (a) Variables were measured on up to 14860 individual plants from 603 species. (b) Coverage of the dataset across global climate space. Grey hexagons indicate the number of 0.5° cells with woody vegetation across the space. Colour symbols show the locations of sampled individuals for three dominant woody functional types.

A second objective was to study relationships between biomass partitioning and climate variables or biome at a global scale. The dataset includes observations of biomass and size metrics for individual plants, compiled from 175 studies across nine vegetation types (Fig. S1), across the three major biomes (boreal, temperate and tropical). In this paper we focus on field-grown woody plants, spanning the entire size range of woody plants (0.01 - *>*100 m height). One key challenge is thus to account for the very large size variation commonly found in any allometric variable (Niklas, 1994). We do this by fitting a semi-parametric data-driven statistical model, i.e. one that does not assume a particular functional form, thereby allowing us to study PFT patterns at a common plant height.

## Materials and Methods

### Terminology

The terms ‘partitioning’ and ‘allocation’ have been used in various ways, confusing comparisons between studies (Litton *et al.*, 2007). Here, we define biomass partitioning as the actual distribution of biomass between compartments (e.g., leaves vs. stems), and biomass allocation as the proportion of net primary production (NPP) that is allocated to some compartment. The two processes are different because of continuous turnover of biomass, which differs strongly between compartments. We may write (McMurtrie & Wolf, 1983),

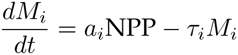

where *M*_*i*_ is the biomass in some compartment (leaves, stems or roots) remaining on the plant, *a*_*i*_ the annual allocation of NPP to that compartment, and *τ*_*i*_ the annual turnover (or loss) of compartment *i* from the plant. It is thus easy to see that partitioning (*M*_*i*_*/M*_*T*_, where *M*_*T*_ is total biomass) can be different from allocation (*a*_*i*_) because turnover (*τ*_*i*_) differs between leaf and wood biomass. Here, we present data on biomass partitioning, which can inform models of allocation only when estimates of turnover are available.

### Data

We used the recently compiled Biomass And Allometry Database (BAAD) (Falster *et al.*, 2015), which in total includes records for 21084 individuals. The database has very limited overlap (n = 261, 1.7 %) with the recent large compilation of Poorter *et al.* (2015) and differs in that measurements are all for individual plants (where Poorter *et al.* (2015) included many stand-based estimates converted back to invididuals). In this paper we restrict our analysis to records that include leaf mass (*M*_*F*_), leaf area (*A*_*F*_), above-ground woody biomass (*M*_*S*_), plant height (*H*), and stem area measured at ground level (*A*_*S*_), or at breast height (typically 1.3m) (*A*_*Sbh*_) (n=14860). The database contains many more variables, for example root biomass for a much smaller subset of studies. Here we limit the analysis to patterns in aboveground biomass distribution. For each analysis, we used different subsets of the data because not all variables were measured in each study. Sample sizes by PFT are summarised in Table 1. We excluded glasshouse and common garden studies, including only field-grown woody plants (including natural vegetation, unmanaged and managed plantations). We considered three PFTs : evergreen angiosperms, evergreen gymnosperms, and deciduous angiosperms. We excluded deciduous gymnosperms because few data were available. All locations were further separated into boreal (including sub-boreal), temperate, and tropical biomes. To assess the coverage of the global climate space by the dataset, we extracted mean annual temperature and precipitation from Worldclim (Hijmans *et al.*, 2005), excluding areas without woody vegetation (taken from the global land cover database GLC-SHARE (Latham *et al.*, 2014)).

**Table 1.** Sample sizes used in the analyses of the four variables considered in the study by plant functional type (PFT, DA = deciduous angiosperm, EA = evergreen angiosperm, EG = evergreen gymnosperm) and biome. Sample sizes denote number of individuals, with number of unique species in parentheses.

| Variable | PFT | boreal | temperate | tropical | Sum |
| --- | --- | --- | --- | --- | --- |
| $M_F/M_T$ | DA | 184 (4) | 1975 (94) | 162 (14) | 2321 (112) |
|  | EA |  | 1271 (68) | 1674 (169) | 2945 (237) |
|  | EG | 669 (7) | 1222 (24) |  | 1891 (31) |
|  | Sum | 853 (11) | 4468 (186) | 1836 (183) | 7157 (380) |
| $A_F/M_T$ | DA | | 1452 (68) | 35 (7) | 1487 (75) |
|  | EA |  | 1124 (59) | 1320 (112) | 2444 (171) |
|  | EG | 321 (7) | 573 (15) |  | 894 (22) |
|  | Sum | 321 (7) | 3149 (142) | 1355 (119) | 4825 (268) |
| $M_F/A_S$ | DA | 308 (6) | 2067 (93) | 162 (14) | 2537 (113) |
|  | EA |  | 1094 (59) | 1696 (173) | 2790 (232) |
|  | EG | 1010 (10) | 1896 (31) |  | 2906 (41) |
|  | Sum | 1318 (16) | 5057 (183) | 1858 (187) | 8233 (386) |
| $A_F/A_S$ | DA | | 1572 (70) | 35 (5) | 1607 (75) |
|  | EA |  | 970 (54) | 1370 (120) | 2340 (174) |
|  | EG | 458 (8) | 1139 (21) |  | 1597 (29) |
|  | Sum | 458 (8) | 3681 (145) | 1405 (125) | 5544 (278) |

For the above-ground biomass pools, we calculated the leaf mass fraction (*M*_*F*_ */M*_*T*_, where *M*_*T*_ is total above-ground biomass) and leaf area ratio (*A*_*F*_ */M*_*T*_). These variables are related by,

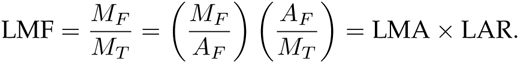

We only used LMA directly estimated for the harvested plants (typically for a subsample of leaves, see Falster *et al.* (2015) for details on the methods for each contributed study). For conifers, leaf area was converted to half-total surface area using the average of a set of published conversion factors (Barclay & Goodman, 2000), with different conversion factors applied to pines (*Pinus* spp.) and non-pines. This conversion was necessary because half-total surface area is most appropriate for comparison to flat leaves (Lang, 1991; Chen & Black, 1992). Stem cross-sectional area was measured either at breast height and/or at the base of the plant. For the subset of the data where both were measured, we estimated basal stem area (*A*_*S*_) from breast height stem area (*A*_*Sbh*_) using a non-linear regression model, as follows.

Using the subset of data where basal stem diameter (*A*_*S*_) and diameter at breast height (*A*_*Sbh*_) were measured (a total of 1270 observations covering the three major PFTs), we developed a non-linear model to predict *A*_*S*_ when only *A*_*Sbh*_ and plant height (*H*) were measured. We fit the following equation, using non-linear regression.

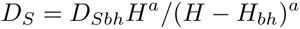

where *D*_*S*_ is the basal stem diameter (m), *D*_*Sbh*_ stem diameter at breast height, *H*_*bh*_ the height at which *D*_*Sbh*_ was measured (typically 1.3 or 1.34m), and *a* was further expressed as a function of plant height:

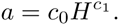

The estimated coefficients were *c*0 = 0.424, *c*1 = 0.719, root-mean square error = 0.0287.

### Data analysis

We used generalised additive models (GAM) to capture the relationships between biomass and plant size variables, and to estimate variables and their confidence intervals such as LMF at a common plant height. In all fitted GAMs, we used a cubic regression spline. For the smoothed term in the model (plant height), we used up to 3 or 4 degrees of freedom, which resulted in biologically realistic smoothed relationships. Within the GAM, we used a penalised regression smoother (Wood, 2006) to allow the final degree of smoothness to be estimated from the data. In all fitted GAMs, we used species-dataset combination as a random effect. All variables (except MAP and MAT) were log-transformed prior to analysis.

Variance explained by quantitative climate variables (MAP and MAT) were tested with GAMs where variables were sequentially added to the model, and the explained variance (*R*^2^) calculated. For the test of biome effects on biomass partitioning, we used a linear mixed-effects model (because two factors and their interactions were tested), again with species-dataset as the random effect. We calculated the *R*^2^ for linear mixed-effects models for the fixed effects only (Nakagawa & Schielzeth, 2013).

Despite the exceptional size of our dataset, the strong size-dependence of LMF still hinders comparisons across climatic gradients, due to the small sample size available within each species or site when sampling at a common height. To study climate effects on biomass partitioning, we therefore further decomposed LMF as the product,

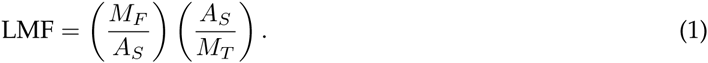

This decomposition showed that the vast majority of size-related variation was captured by *A*_*S*_ */M*_*T*_ alone, indicating that comparisons among *M*_*F*_ */A*_*S*_ could be made without needing to compare at a common height. A similar decomposition of LAR as the product of *A*_*F*_ */A*_*S*_ and *A*_*S*_ */M*_*T*_ produced the same outcome. We therefore analysed for climatic effects on *M*_*F*_ */A*_*S*_ and *A*_*F*_ */A*_*S*_ in two ways: with quantitative variables (mean annual precipitation, MAP; mean annual temperature, MAT), and by biome (tropical / temperate/ boreal), a simple classification taking into account both MAT and MAP (Fig. S1). In both cases we used PFT and plant height as covariates varying across climate space.

All analyses were conducted in R v3.2.0 (R Core Team, 2015). GAMs were fitted using *mgcv* package (Wood, 2006). The code to replicate this analysis are available on GitHub at http://github.com/RemkoDuursma/baadan

## Results

The raw data in Fig. 2 show a steeper increase of woody aboveground biomass (*M*_*S*_) with plant height compared to foliage biomass (*M*_*F*_). As a result, LMF decreased with plant height, with the three plant functional types clearly differing in LMF across nearly the entire size range (Fig. 3a). When we corrected for plant height by estimating LMF at a common plant height, we found that LMF was proportional to the average leaf mass per area (LMA) across the three PFTs (Fig. 3b). As a consequence, LAR was invariant between PFTs, because LAR = LMF / LMA (Fig. 3c and Fig. S2).

**Fig. 2.**
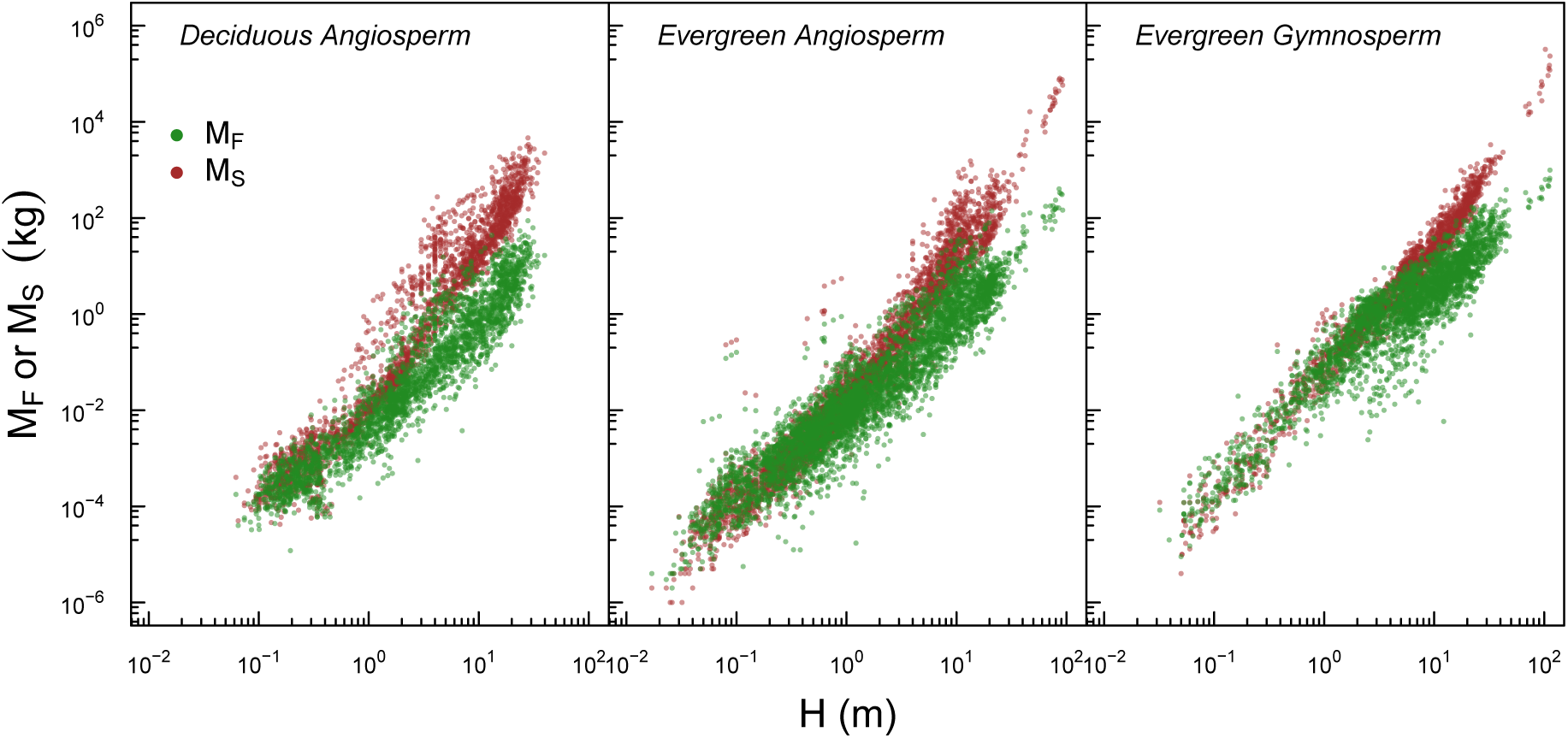
Raw data for leaf biomass (*M*_*F*_) and total above-ground woody biomass (*M*_*S*_) for each of the PFTs, as a function of total plant height (*H*). Each point is an individual plant. Sample sizes are listed in Table 1

Mirroring the results for LMF, we found that the amount of leaf mass per unit stem area (*M*_*F*_ */A*_*S*_) differed among the three PFTs, that these differences were also proportional to LMA (Fig. 3d), and that PFTs were similarly invariant in the amount of leaf area per unit stem area (*A*_*F*_ */A*_*S*_) (Fig. 3e).

**Fig. 3.**
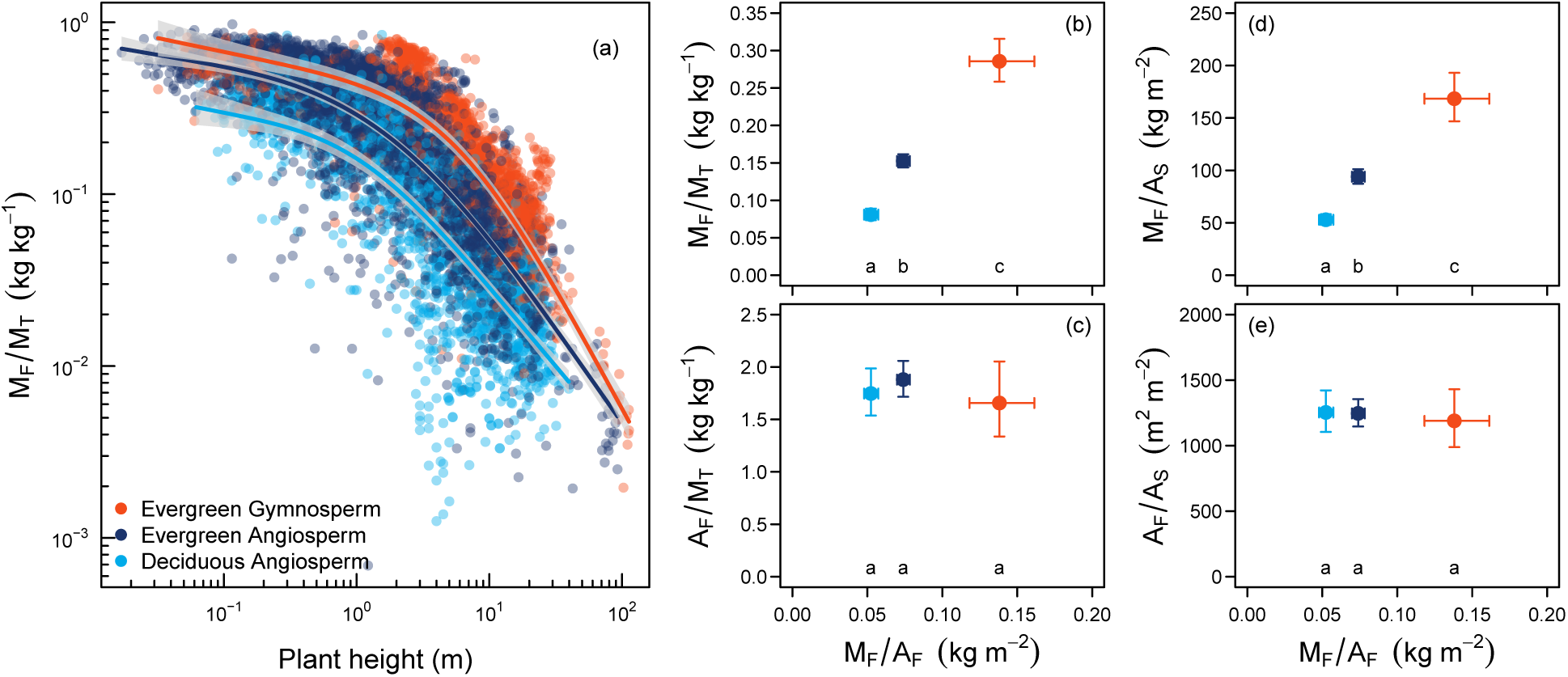
Dominant woody PFTs differ in leaf mass fraction due to underlying differences in leaf mass per area. (a) Leaf mass fraction (*M*_*F*_ */M*_*T*_ = leaf mass / above-ground biomass) by PFT. Each symbol is an individual plant. Lines are generalised additive model fits. (b) and (c) Leaf mass fraction and leaf area ratio (*A*_*F*_ */M*_*T*_) at the average plant height in the dataset, estimated from panel (a). (d) Average leaf mass per unit basal stem area, and (e) leaf area per unit basal stem area for the three major PFTs confirm that the between-PFT variation in leaf mass fraction is due to leaf mass per unit basal area. Error bars are 95% confidence intervals. Letters denote significant differences (at *α* = 0.05).

The above results highlight how LMA drives differences in biomass partitioning among dominant woody PFTs. To assess whether LMA could replace PFT when predicting biomass partitioning, we fit a linear mixed-effects model to LMF and *M*_*F*_ */A*_*S*_ using PFT and plant height (and the quadratic term) as predictors (and all interactions). We then replaced PFT with LMA, and found that the model with LMA could explain almost as much variation in LMF as the model with PFT (*R*^2^ = 0.74 with PFT vs. 0.62 with LMA), likewise for *M*_*F*_ */A*_*S*_ (*R*^2^ = 0.29 vs. 0.28).

There was considerable variation in all studied variables between species within PFTs (Fig. 4). To understand whether we can improve on a PFT-based classification by including other traits that affect biomass partitioning, we decomposed *M*_*F*_ */A*_*S*_ into LMA (*M*_*F*_ */A*_*F*_), and the ratio of leaf area to stem cross-sectional area (*A*_*F*_ */A*_*S*_):

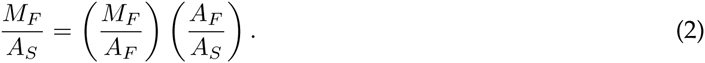

**Fig. 4.**
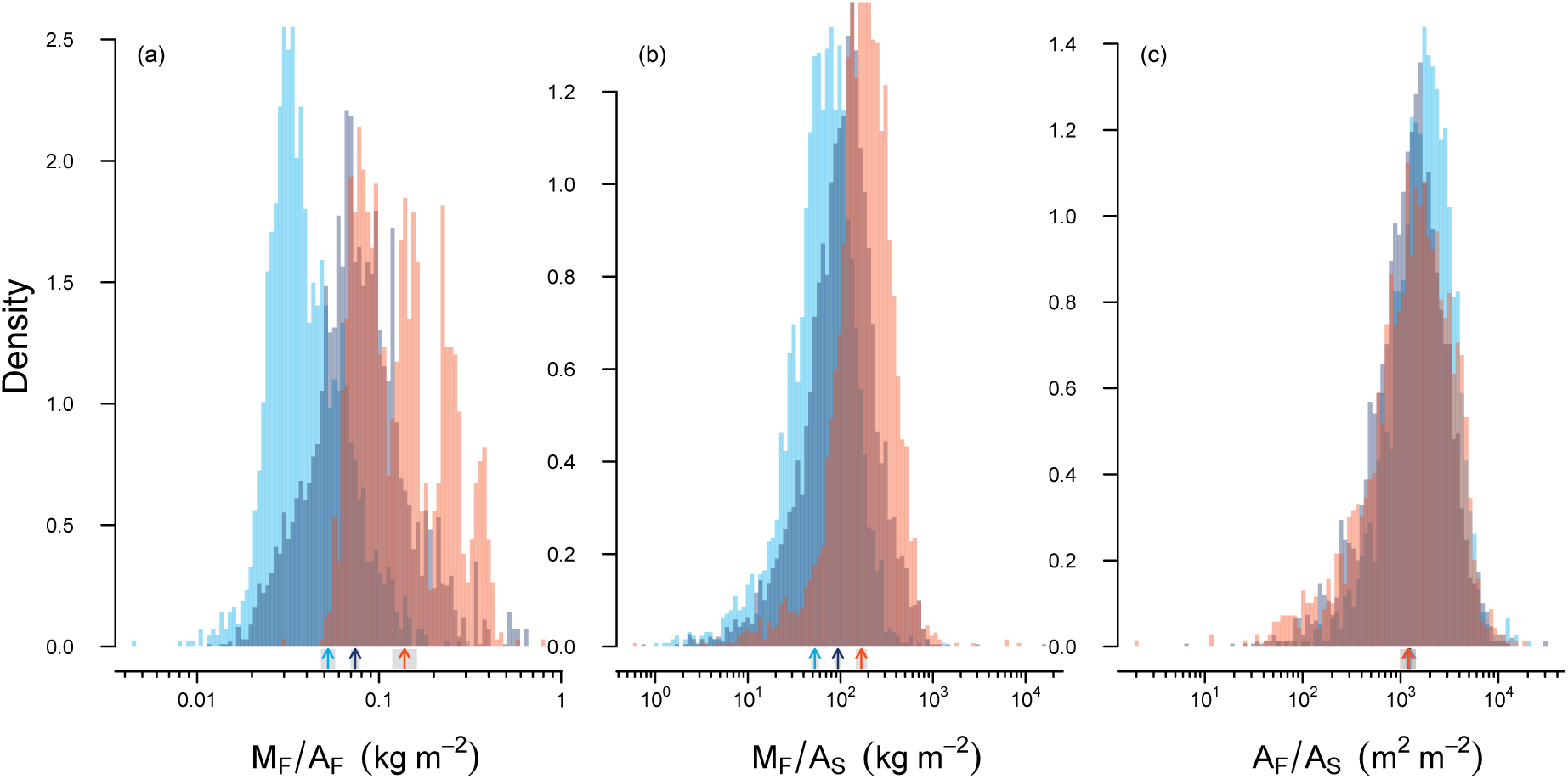
Plant functional types diverge strongly in mass-based partitioning, but converge in area-based partitioning. Shown are histograms (as probability density functions) of leaf mass per area (*M*_*F*_ */A*_*F*_), leaf mass per unit basal stem area (*M*_*F*_ */A*_*S*_) and leaf area per unit basal stem area (*A*_*F*_ */M*_*S*_) grouped by the three PFTs. Arrows indicate means by PFT. Colours as in Fig. 3.

The ratio *M*_*F*_ */A*_*S*_ is relevant because plant leaf mass is frequently estimated from *A*_*S*_ with allometric equations (Shinozaki *et al.*, 1964; Chave *et al.*, 2005), using records of stem diameter commonly recorded on long-term monitoring plots. The correlation between *M*_*F*_ */A*_*S*_ and LMA was found to hold also within PFTs (Fig. 5a), explaining 30% of the variation in *M*_*F*_ */A*_*S*_ across species. The regressions across species by PFT were broad and overlapping, with a more general relationship extending across the entire LMA axis. A larger fraction of the variation in *M*_*F*_ */A*_*S*_ was explained by species-level differences in *A*_*F*_ */A*_*S*_ (Fig. 5b). As previously noted, *A*_*F*_ */A*_*S*_ does not differ systematically among PFTs (Fig. 3e), but does vary close to two orders of magnitude across species. In Fig. 5b, the separation among PFTs in *M*_*F*_ */A*_*S*_ arises due to differences in LMA.

**Fig. 5.**
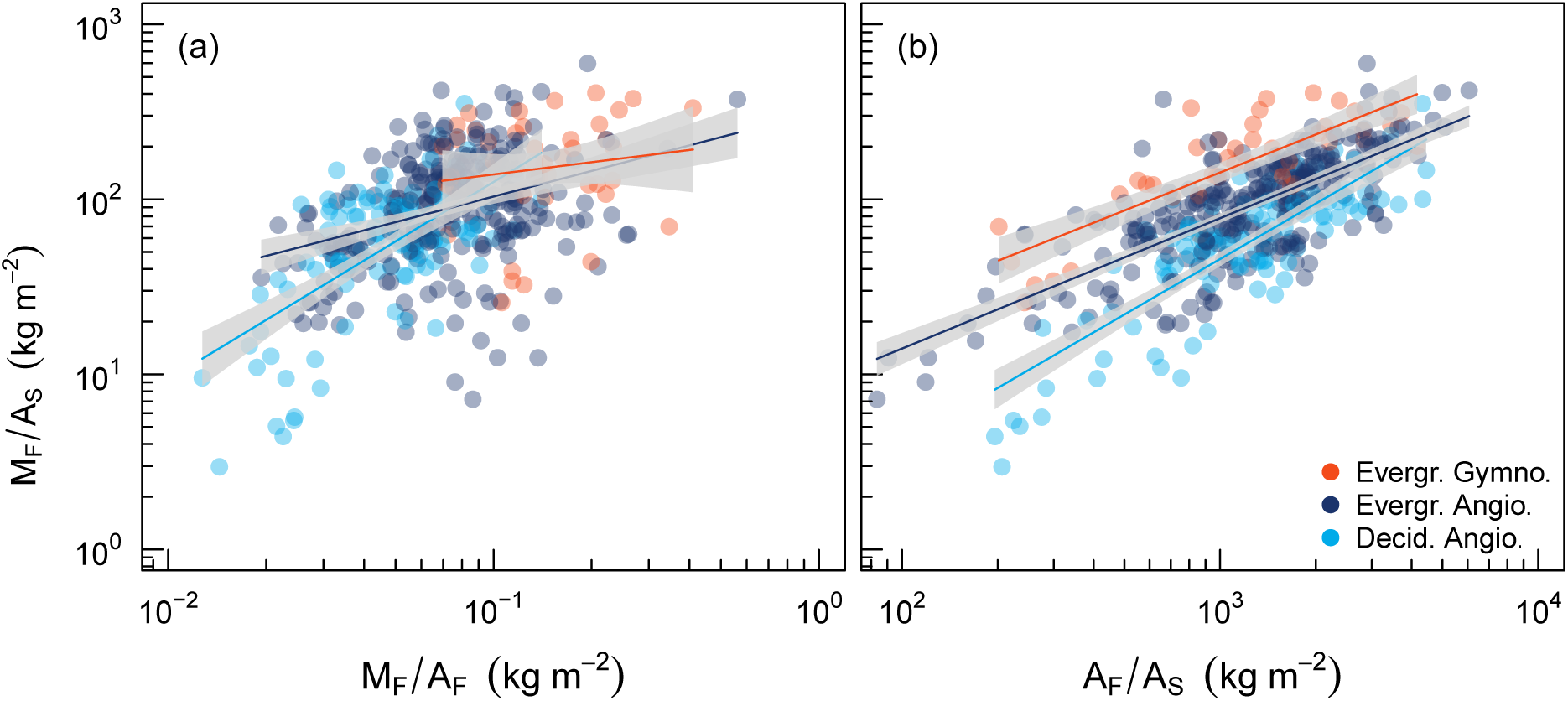
Leaf and stem traits capture variation in biomass partitioning across species. Individual plant data were averaged by species and study combinations. Lines are linear regressions with 95% confidence intervals. Both regressions were significant (*P <*0.01). *R*^2^ for fitted relationships are 30.9% in panel (a) and 66% in panel (b) (for a linear regression model including PFT as a factor variable).

We found that decomposing LMF and LAR as shown in Eq. 1 was very useful to study climate effects because the second term (*A*_*S*_ */M*_*T*_) absorbed nearly all of the size-related variation in LMF (Table 2 and Fig. S3) and is otherwise fairly conserved across PFTs (Fig. S3). As shown in Fig. 5b and Fig. 3d, the term *M*_*F*_ */A*_*S*_ exhibits comparable differences between PFTs to LMF, but unlike LMF, *M*_*F*_ */A*_*S*_ is nearly independent of plant height (Table 2). *M*_*F*_ */A*_*S*_ is thus a useful proxy for LMF that can be compared across species and sites.

**Table 2.** Explained variance by plant functional type (PFT), plant height (H) and climate variables (either biome (B), or MAP and MAT) for five whole-plant variables. For the analysis with biome, each variable was added to a linear mixed-effects model, using species within dataset as a random effect. The *R*^2^ shown is that explained by the fixed effects only. All fixed effects were highly significant (P *<*0.001). In addition to the fixed effects shown, all interactions were added to each of the models. For the analysis with continuous climate variables MAP and MAT, each variable was added as a smooth term in a generalized additive model (GAM), with the exception of PFT (a categorical variable). Note that explained variance for the linear mixed-effects model and GAM is not necessarily the same even with the same predictors, due to different methods for fitting and estimating the explained variance.

| Mixed-effects model |  | Predictors |  |  |
| --- | --- | --- | --- | --- |
| Variable | H | H, PFT | H, PFT, B |  |
| Leaf mass fraction($M_F/M_T$) | 0.65 | 0.75 | 0.76 | |
| Leaf area ratio($A_F/M_T$) | 0.69 | 0.69 | 0.75 | |
| Leaf mass / stem basal area($M_F/A_S$) | 0.01 | 0.25 | 0.29 | |
| Leaf area / stem basal area($A_F/A_S$) | 0.01 | 0.03 | 0.05 | |
| Leaf mass per area($M_F/A_F$) | 0.13 | 0.60 | 0.66 | |
| GAM |  | Predictors |  |  |
| Variable | H | H, PFT | H, PFT, MAT, MAP |  |
| Leaf mass fraction($M_F/M_T$) | 0.65 | 0.76 | 0.74 | |
| Leaf area ratio($A_F/M_T$) | 0.69 | 0.71 | 0.73 | |
| Leaf mass / stem basal area($M_F/A_S$) | 0.03 | 0.24 | 0.25 | |
| Leaf area / stem basal area($A_F/A_S$) | 0.05 | 0.12 | 0.23 | |
| Leaf mass per area($M_F/A_F$) | 0.12 | 0.47 | 0.61 | |

Regardless of how we analysed the data, we found only weak climatic effects on biomass partitioning within PFTs (Fig. 6). Biome consistently explained very little variation when added to a statistical model in addition to PFT and plant height (*R*^2^ increased by only 0.01 - 0.06, see Table 2). Likewise, we found weak and inconsistent effects of climate variables (MAP and MAT) (Fig. 6b-c). These variables explained little variation when added to a statistical model in addition to PFT and plant height (*R*^2^ increased by 0.01 and 0.11, respectively, see Table 2).

**Fig. 6.**
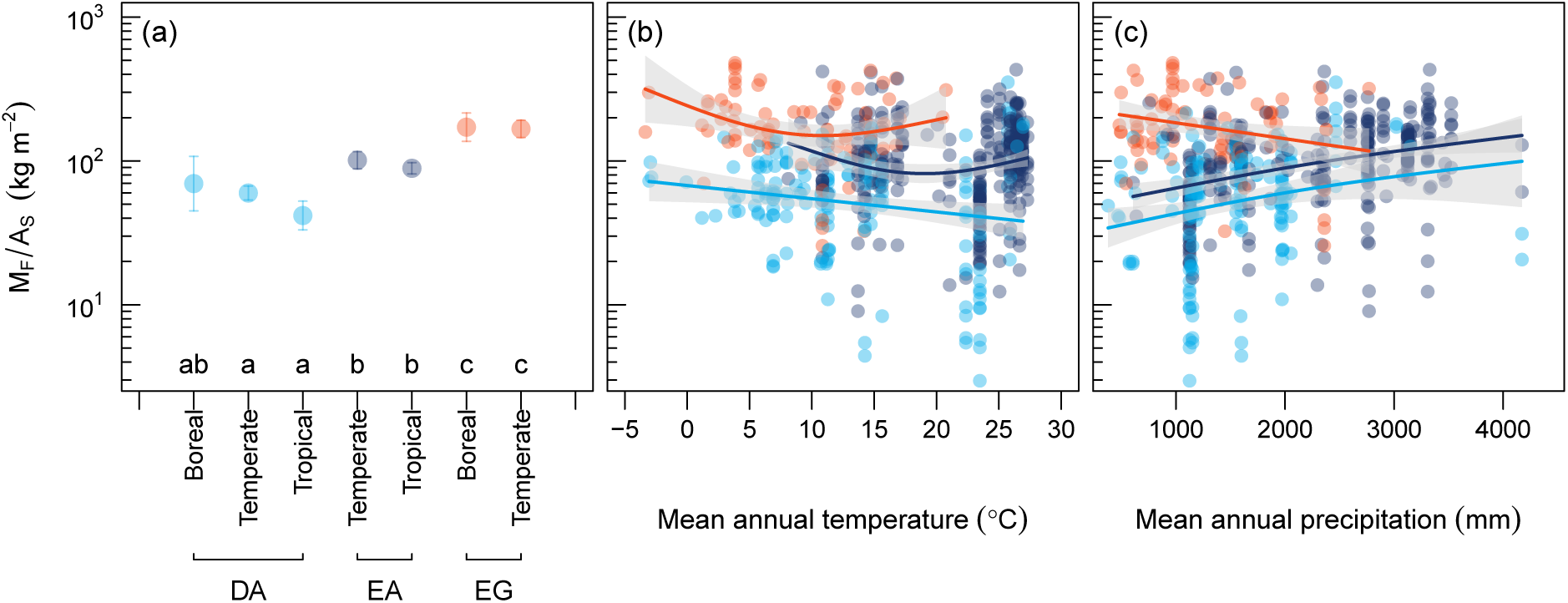
Beyond influencing the distribution of PFTs, climate only modestly influences biomass partitioning. a) Leaf mass per unit stem cross-sectional area (*M*_*F*_ */A*_*S*_) for each PFT within three different biomes. DA: deciduous angiosperms, EA: evergreen angiosperms, EG: evergreen gymnosperms. Error bars are 95% confidence intervals for the mean, estimated from a linear mixed-effects model using species within study as the random effect. Letters denote significant differences (at *α* = 0.05). b-c) Relationships between climate and *M*_*F*_ */A*_*S*_ within each PFT. Solid lines are generalised additive model (GAM) fits (with a base dimension of 3); with grey areas indicating the 95% confidence interval around theGAM.

## Discussion

We found that PFTs differed in LMF, and that LMF was proportional to LMA. This conclusion was particularly strong across the three PFTs studied, but it also held across species within PFT. The implication is that the amount of leaf area supported per unit biomass (leaf area ratio, LAR) does not differ between PFTs. This result seems robust, as our data include individual plants spanning the entire size range of woody plants in natural forests (0.1-100m) (Fig. 3a). Previous studies have demonstrated a large difference in LMF between angiosperms and gymnosperms (Poorter *et al.*, 2012, 2015), but have not been able to explain these differences in terms of leaf traits. We show that LMA – a central trait of the leaf economics spectrum (LES) (Wright *et al.*, 2004) – explains differences in LMF in a consistent manner.

We found that, as expected, plant size strongly influences LMF (Fig. 3) and LAR (Fig. S2) (see also Poorter *et al.* (2012, 2015)). It is thus necessary to correct for plant size when comparing biomass partitioning parameters, as has been noted many times (McConnaughay & Coleman, 1999). We used a semi-parametric approach to account for plant size, which has the advantage that it does not require an *a priori* assumption on the functional relationship. This was useful because both LMF and LAR showed very non-linear patterns with plant height, even on a logarithmic scale, consistent with recent other results (Poorter *et al.*, 2015).

The finding that LMF is proportional to LMA across PFTs implies that LAR does not differ systematically between PFTs. One explanation for this pattern lies in the strong positive correlation between LMA and leaf lifespan (LL) (Wright *et al.*, 2004). Both Poorter *et al.* (2012) and Enquist & Niklas (2002) explained higher LMF in gymnosperms by higher LL; plants simply maintain more cohorts of foliage, but this explanation demands that all else is equal. Yet it is not self-evident that everything else is indeed equal, that is, annual production of foliage could differ between PFTs. Nonetheless, further supporting this explanation, Reich *et al.* (1992) showed a positive correlation between LL and total stand leaf biomass across diverse forest stands. Another, not mutually exclusive, possible explanation is that plants may allocate biomass in a manner that targets a more constant LAR rather than LMF. Canopy size scales with *M*_*T*_ (Duursma *et al.*, 2010), and a larger canopy means higher light interception per unit total biomass (Duursma & Mäkelä, 2007). It can thus be argued that an optimal LAR exists that balances investment in supporting woody biomass (reflected by *M*_*T*_) and the efficiency of foliage in terms of light interception.

We found that patterns in *M*_*F*_ */A*_*S*_ across PFTs mirrored those of LMF (Fig. 3). Because LMF may be decomposed into the terms *M*_*F*_ */A*_*S*_ and *M*_*T*_ */A*_*S*_ (eq. 2), this result suggests that *M*_*T*_ */A*_*S*_ does not vary between PFTs (see also Fig. S3). If this result holds, it means that aboveground biomass may be estimated across PFTs with a single equation that does not differ between PFTs. For tropical forests, Chave *et al.* (2005) showed that aboveground biomass can be estimated as,

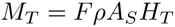

where *F* is a form factor, and *ρ* wood density (kg m^3^). After rearranging, this equation predicts that *M*_*T*_ */A*_*S*_ is proportional to plant height. We found some support for this prediction (Fig. S3), suggesting further improvements could be made by incorporating this dependence. It is timely to use database made public by Falster *et al.* (2015) (BAAD) and Poorter *et al.* (2015) to develop more general equations to estimate individual plant aboveground biomass across the globe.

We found that climate did not appreciably affect biomass partitioning between leaves and stems. This finding is in contrast with a recent study where LMF was found to increase with MAT across the globe (Reich *et al.*, 2014). One explanation for this difference is that our measurements were taken on individual plants, whereas in Reich *et al.* (2014) they were on whole stands. This suggests climatic effects on stand-level biomass partitioning occur primarily by altering the size-distribution and stand density (number of individuals per unit area), rather than partitioning within individual plants. With the caveats that there is still variation between species and studies unaccounted for (Fig. 4), and that coarse climate variables such as MAT and MAP may mask small-scale climate effects on biomass partitioning, our results strongly suggest the effects of PFT are stronger than any direct climate effect on biomass partitioning within individual plants (Table 2).

Our results indicate that it is possible to integrate a key leaf trait with whole-plant modelling of biomass partitioning. Indeed, nearly as much variation was explained across species when using LMA instead of PFT in a statistical model. This is surprising because LMA only captures one aspect of functional differentiation among PFT – leaf morphology (but including correlated effects on leaf lifespan and photosynthetic capacity via the leaf economics spectrum (Wright *et al.*, 2004)). We thus show that in a PFT-based classification, LMA is a good first estimate of biomass partitioning, however, more variation can be explained by another trait (*A*_*F*_ */A*_*S*_), which varies appreciably between species. These results suggest promising avenues for parameterising and simplifying biomass partitioning routines in GVMs.

Overall, our results establish general patterns about plant construction and thus lay an empirical base against which models can be benchmarked. A recent study compared allocation routines in a number of leading ecosystem models (De Kauwe *et al.*, 2014), and recommended constraining allocation by observed biomass fractions instead of using constant allocation fractions. Based on our findings, a first approximation within GVMs would be to assume leaf area to stem cross-sectional area a parameter that does not vary between PFTs (notwithstanding effects of plant height on this variable), and vary biomass allocation accordingly. LMA, already a parameter in most GVMs, then gives the ratio of leaf biomass to stem area. Some models already incorporate a similar algorithm (see Notes S1), while other algorithms may be tuned to yield similar patterns between PFTs as we have presented. In any case, the growing availability of large datasets on stand biomass and individual plant construction (Falster *et al.*, 2015) suggest the time is ripe for rigorous benchmarking (Abramowitz, 2012; De Kauwe *et al.*, 2014) of GVMs against empirical data.

## Acknowledgments

Our sincere thanks to everyone who contributed data to the Biomass and Allometry Database. We thank Martin De Kauwe and anonymous referees for comments on an earlier version of this paper. Thanks also to Rich FitzJohn for advice about using the remake package. Figure 2a uses an image by Lana Heydon, IAN Image Library http://ian.umces.edu/imagelibrary.

## Supporting Information

Additional supporting information may be found in the online version of this article.

Fig. S1 Global coverage of the climate space by the dataset, labelled by vegetation type.

Fig. S2 Leaf area ratio (*A*_*F*_ */M*_*T*_) by PFT.

Fig. S3 Relationship between above-ground biomass and basal stem area.

**Notes S1** Modelling of biomass partitioning in global vegetation models (GVMs))

